# Deregulation of the histone H3K9 di-methylation landscape suppresses canonical Wnt signaling in embryonal rhabdomyosarcoma

**DOI:** 10.1101/2020.04.20.050120

**Authors:** Ananya Pal, Jia Yu Leung, Gareth Chin Khye Ang, Vinay Kumar Rao, Luca Pignata, Huey Jin Lim, Maxime Hebrard, Kenneth T Chang, Victor KM Lee, Ernesto Guccione, Reshma Taneja

## Abstract

The Wnt signaling pathway is down-regulated in embryonal rhabdomyosarcoma (ERMS) and contributes to the block of myogenic differentiation. Epigenetic mechanisms leading to its suppression are unknown and could pave the way towards novel therapeutic modalities. In this study, we demonstrate that the H3K9 lysine methyltransferase G9a suppresses canonical Wnt signaling by activating expression of the Wnt antagonist *DKK1*. Inhibition of G9a expression or activity reduced *DKK1* expression and elevated canonical Wnt signaling resulting in myogenic differentiation *in vitro* and *in vivo*. Mechanistically, G9a impacted Sp1 and p300 enrichment at the *DKK1* promoter in a methylation-dependent manner. The reduced tumor growth upon G9a deficiency was reversed by recombinant DKK1 or LGK974, which also inhibits Wnt signaling. Consistently, among thirteen drugs targeting chromatin modifiers, G9a inhibitors were highly effective in reducing ERMS cell viability. Together, our study demonstrates that ERMS cells are vulnerable to G9a inhibitors and suggest that targeting the G9a-DKK1-β-catenin node holds promise for differentiation therapy.

## Introduction

Rhabdomyosarcoma (RMS) is the most common malignant soft tissue sarcoma (1–3) that arises due to a block in myogenic differentiation. Children with high risk disease have poor prognosis with only 30% showing 5-year event free survival. Embryonal rhabdomyosarcoma (ERMS) accounts for the majority (~60%) of all RMS cases. No single genetic lesion is linked to ERMS but chromosome gains (chr 2,8, 12 and 13) and loss of heterozygosity at 11p15.5 are characteristically seen (4). A few recurrent mutations occur in ERMS that include mutations in p53 (TP53), amplification of CDK4, upregulation of MYCN, and point mutations in RAS leading to its activation (1, 4–6). Recent studies have investigated whether improper epigenetic imprinting underlies the myogenic differentiation defect in RMS (7). This includes altered expression of histone deacetylases, methyltransferases as well as lncRNAs and microRNAs that inhibit differentiation. Among these, EZH2 which mediates repressive histone H3 lysine 27 trimethylation (H3K27me3) is upregulated and binds to muscle specific genes in ERMS. Its silencing increases both MyoD binding and transcription of target genes (8). Similarly, HDAC inhibitors have been found to induce differentiation and reduce self-renewal and migratory capacity of ERMS by regulating Notch-1 and EphrinB1 mediated pathways (9). Interestingly, overexpression of the lysine methyltransferase SUV39H1 suppresses tumor formation in KRAS^G12D^-driven zebrafish model of ERMS (10).

Genetic models that develop ERMS-like tumors due to deregulation of key signaling pathways such as Hedgehog, Wnt, Notch and YAP signaling have been described (11). Double mutants lacking p53 and c-fos (p53/-/c-fos-/-) develop ERMS. Elevated expression of Wnt antagonists dickkopf-related protein 1 (DKK1) and secreted frizzled-related proteins (sFRPs), as well as downregulation of Wnt agonists such as Wnt ligands Wnt 7b, Wnt 5a, Wnt 4 and Wnt 11 was reported in these tumors (12). Mice expressing activated smoothened under the AP2-Cre promoter leading to activation of Hedgehog signaling also develop ERMS. The tumors also show upregulation of DKK3 which similar to DKK1, inhibits canonical Wnt signaling (13). Consistent with these findings, GSK3β inhibitors that activate Wnt signaling were most effective inducers of differentiation in a zebrafish model of ERMS (14). Importantly down regulation of Wnt signaling was found to be relevant only in ERMS.

There are 19 Wnt ligands, which function either through the canonical or non-canonical pathways in a highly context-dependent manner (15). Canonical signaling is activated when Wnt ligands bind to a receptor from the Frizzled (Fzd) family. The co-receptors LRP5/6 facilitate Wnt signaling. Activation of Wnt signaling leads to disruption of the destruction complex [APC, Axin, GSK3β and CK1α] which phosphorylates and degrades β-catenin. Induction of Wnt signaling results in accumulation of non-phosphorylated β-catenin (active β-catenin). β-catenin then translocates to the nucleus where it activates genes in cooperation with TCF/LEF1, but also other transcription factors (15). Neither of the two non-canonical pathways [Planar Cell Polarity pathway (PCP) and the Wnt/calcium signaling pathway] involve β-catenin. DKK1, a secreted protein interacts with LRP5/6 and antagonizes Wnt signaling by preventing LRP5/6 association with Wnt/Fzd complex (16). Despite the relevance of Wnt signaling in ERMS, epigenetic mechanisms leading to its suppression have not been described and could pave the way to development of targeted therapies.

G9a, a lysine methyltransferase mediates mono and di-methylation of H3K9 (H3K9me1/2), which is primarily involved in transcriptional repression (17). Recent studies however have shown that G9a can also function as an activator in methylation-independent and dependent ways (18, 19). G9a has been proposed to have oncogenic functions and its overexpression in leukaemia, gastric, lung, prostate cancer and alveolar rhabdomyosarcoma causes silencing of tumor suppressor genes through its H3K9me2 activity (18–20). In this study, we found that canonical Wnt/β-catenin signaling is epigenetically suppressed in ERMS. G9a activates expression of *DKK1* in a methylation-dependent manner through an impact on Sp1 and p300 recruitment. Our data indicate the potential of targeting the G9a-DKK1 axis to activate Wnt signaling for the development of novel ERMS therapeutics.

## Results

### G9a inhibitors reduce ERMS cell viability

We recently reported that G9a is overexpressed in ARMS (20). To examine whether G9a expression is de-regulated in ERMS, and if it is functionally relevant in these tumor subtype, we first examined its expression in 16 ERMS patient tumor sections using an anti-G9a antibody. High nuclear expression relative to normal muscle was apparent (Figure 1A). In addition, compared to primary human skeletal muscle myoblasts (HSMM), G9a overexpression at both mRNA and protein levels was apparent in three ERMS patient derived cell lines RD18, JR1 and RD (Figure 1B and Figure 1C). To examine if the G9a pathway is functionally relevant, we treated JR1 and RD cell lines with 13 methyltransferase inhibitors at three different concentrations. Viability was measured 8 days later using MTS assay. Drugs targeting BRD4, PRMT5 and G9a showed a strong effect on viability of both cell lines (Figure 1D and Figure 1E). Consistently, treatment of JR1 cells with UNC0642, a small molecule inhibitor of G9a led to a striking reduction in colony formation (Figure 1F). Together, these results indicate that G9a is overexpressed and functionally relevant in ERMS.

**Figure 1.**
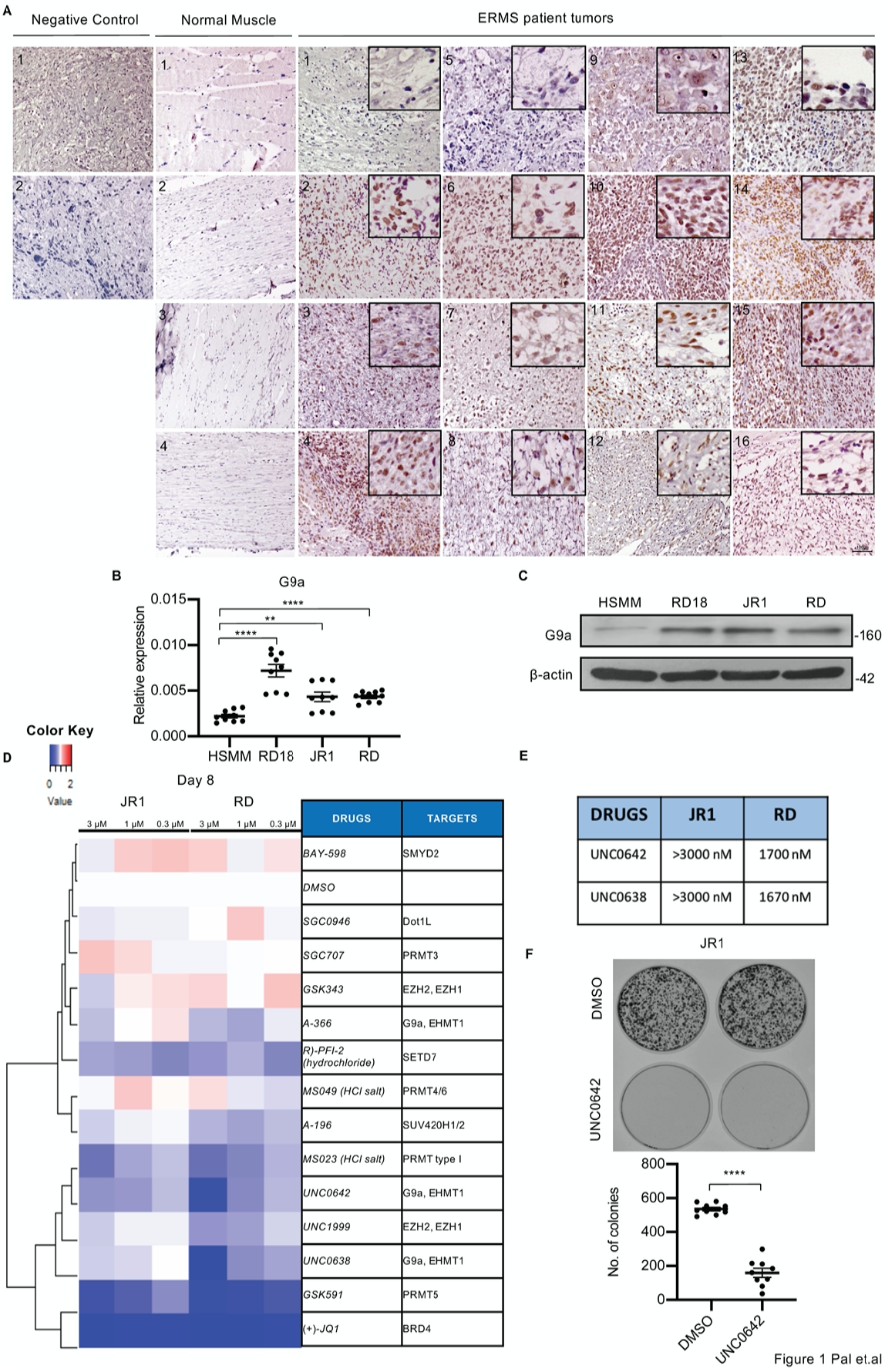
G9a is overexpressed in ERMS. (**A**) 16 archival ERMS patient tumor specimens and 4 normal muscle samples were analysed by immunohistochemistry using anti-G9a antibody. Negative control indicates staining using secondary antibody alone. Inset shows zoomed in image of nuclear G9a staining. Scale bar: 100μm. (**B**) G9a mRNA (n=3) were examined in three patient derived cell lines (RD, RD18 and JR1) in comparison to primary human skeletal muscle myoblasts (HSMM) by qPCR. Values correspond to the average ± SEM. All three ERMS cell lines examined showed an increased G9a mRNA expression compared to HSMM. (**C**) G9a protein levels were examined by western blotting in HSMM, RD18, JR1 and RD cells. A representative image of 3 different experiments is shown. All three ERMS cell lines examined showed an increased G9a protein expression compared to HSMM. (**D**) ERMS cell lines JR1 and RD were treated with the indicated methyltransferase inhibitors (3, 1 and 0.3 μM). Viability at day 8 was scored by MTS assay and measured as the ratio over control cells treated with an equivalent dilution of DMSO. RED indicates viability >control; WHITE is equal to control, and BLUE is less than control. The experiment was conducted in triplicates and (+)-JQ1 was used as a positive control. GSK591, UNC0642, UNC638 and had a strong effect on viability. (**E**) The IC50 of G9a inhibitors UNC0642 and UNC0638 in JR1 and RD cell lines is shown. (**F**) JR1 cells were treated with DMSO or UNC0642 for 10 days. Colony formation was assessed by staining with crystal violet. A representative image of 3 different experiments is shown. The dot plot shows the number of colonies upon DMSO and UNC0642 treatment. Statistical significance in **B** and **F** was calculated by unpaired two-tailed *t* test. \*\**P* ≤ 0.01, \*\*\**P* ≤ 0.001.

### G9a inhibition promotes myogenic differentiation and inhibits proliferation in ERMS cell lines

To examine the role of G9a in ERMS, we depleted its endogenous expression in RD18, JR1 and RD cells using small interfering RNA (Figure 2A, Supplementary Figure S1A, and Supplementary Figure S2A), or blocked its methyltransferase activity using UNC0642 that resulted in reduced H3K9me2 (Figure 2B, Supplementary Figure S1B, and Supplementary Figure S2B). We then examined the impact on differentiation and proliferation of tumor cells. G9a knockdown (siG9a cells), as well as UNC0642 treatment resulted in increased myogenic differentiation relative to their respective controls as evidenced from the increased MHC expression, a terminal differentiation marker, as well as myogenin, an early differentiation marker (Figure 2C, Supplementary Figure S1C and Supplementary Figure S2C). To differentiate, myoblasts irreversibly exit the cell cycle (21). Given the enhanced myogenic differentiation upon G9a depletion, we investigated the impact of G9a loss on proliferation by labelling S-phase cells with BrdU. Both siG9a cells and UNC0642 treatment resulted in a significant decrease in BrdU^+^ cells compared to their respective control in RD18 cells (Figure 2D, Supplementary Figure S1D, and Supplementary Figure S2D). Further, stable G9a knockdown in RD cells also resulted in increased MHC levels and decreased BrdU^+^ cells (Supplementary Figure S2E-G). A striking reduction in colony formation was also seen in shG9a cells compared to controls (Supplementary Figure S2H). Together, these results indicate that G9a inhibition permits cells to exit the cell cycle and undergo myogenic differentiation.

**Figure 2.**
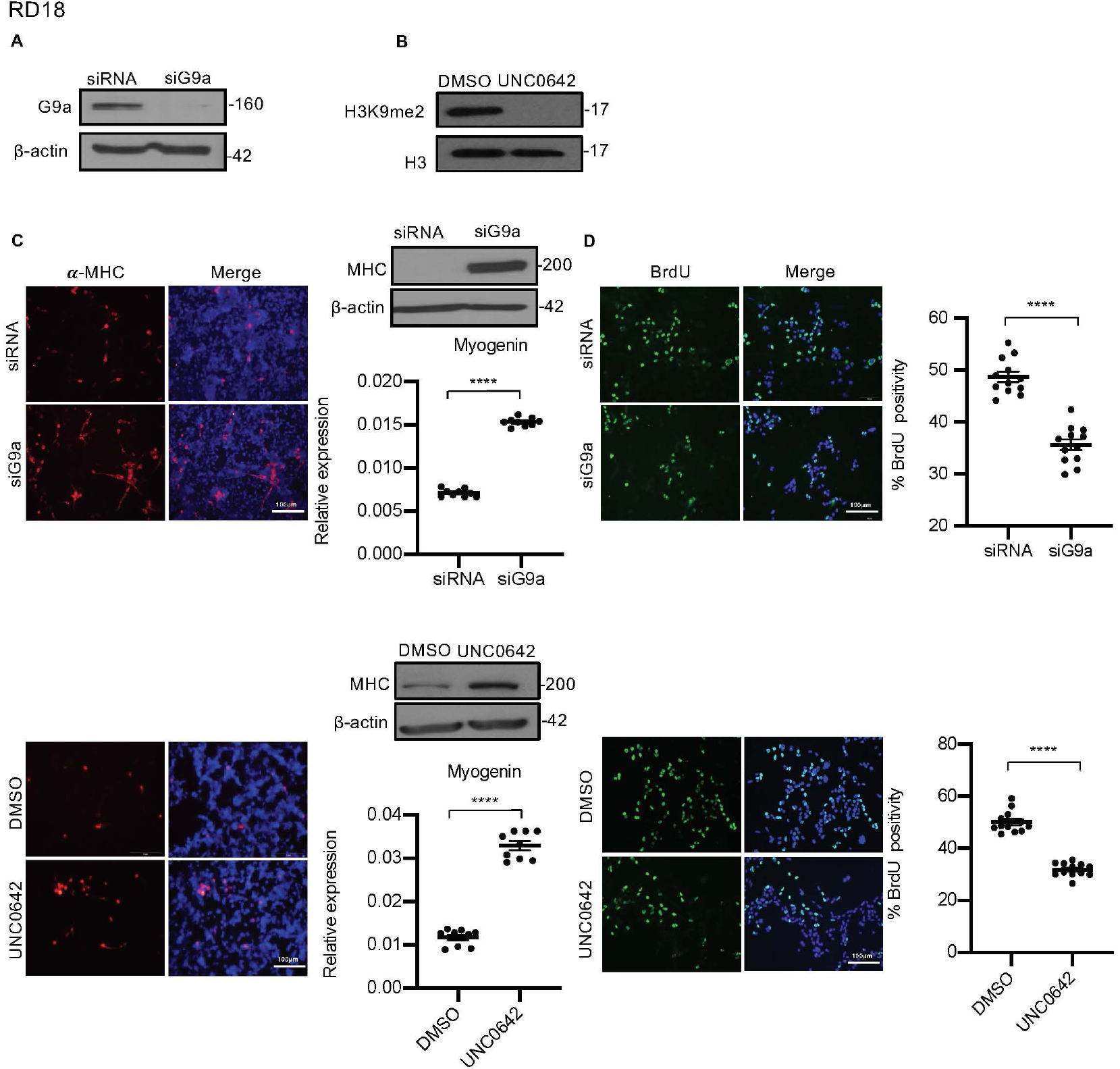
G9a inhibits differentiation and promotes proliferation of myoblasts. (**A**) G9a was depleted in RD18 cells using siRNA. Control and siG9a cells were analysed for knockdown efficiency by western blot. β-actin was used as an internal loading control. (**B**) H3K9me2 levels were analysed 48 hr after 2.5 μM of UNC0642 treatment. Histone H3 was used as a loading control. (**C**) Differentiation was analysed in control and siG9a cells (upper panels) or DMSO and 2.5 μM of UNC0642 treated cells (lower panels) after culture for five days in differentiation medium (DM). Cells were analysed by immunofluorescence and western blot using anti-MHC antibody as indicated. Nuclei were stained with DAPI. Representative images of 3 different experiments are shown. Expression of myogenin was analysed by qPCR at day 2 of differentiation (n=3). Values correspond to the average ± SEM. (**D**) Proliferation was analysed in control and siG9a (upper panels); or DMSO and 2.5 μM of UNC0642 treated cells (lower panels) by immunostaining with anti-BrdU antibody. Cells were analysed by immunofluorescence (n=3). The dot plots show the percentage of BrdU^+^ in siG9a and UNC0642 treated cells relative to their respective controls. Values correspond to the average ± SEM. Statistical significance in **C** and **D** was calculated by unpaired two-tailed *t* test. \*\*\*\**P* ≤ 0.0001.

### G9a regulates DKK1 and canonical Wnt signaling

In order to identify mechanisms underlying G9a function, we performed RNA-Sequencing (RNA-Seq). Cluster analysis of differentially expressed genes from control RD and G9a knockdown cells was done in triplicates (Figure 3A). Volcano plot of differentially expressed genes (Figure 3B) revealed that 872 genes were significantly up regulated in siG9a cells compared to the control, of which 494 genes had a fold change >1.2. Among the 1098 genes that were significantly down regulated in siG9a cells, 695 genes had a fold change >1.2. Gene Ontology (GO) analysis showed that among the top 20 unique biological processes associated with differentially expressed genes in siG9a cells were cell cycle progression and Wnt signaling (Figure 3C). Given its relevance in ERMS, we focused on the Wnt pathway. Interestingly, negative regulators of the Wnt pathway such as DKK1, DKK3 and ITGA3 (22, 23) were downregulated in siG9a cells, whereas positive regulators such as Wnt3 and FRAT2 were upregulated (Supplementary Figure 3A-B). We validated genes in the Wnt pathway (Figure 3D and Supplementary Figure S3C-F) as well as those involved in skeletal muscle differentiation such as MyoD1, Myostatin, MYL and Myozenin by qPCR (Supplementary Figure S3G-J).

**Figure 3.**
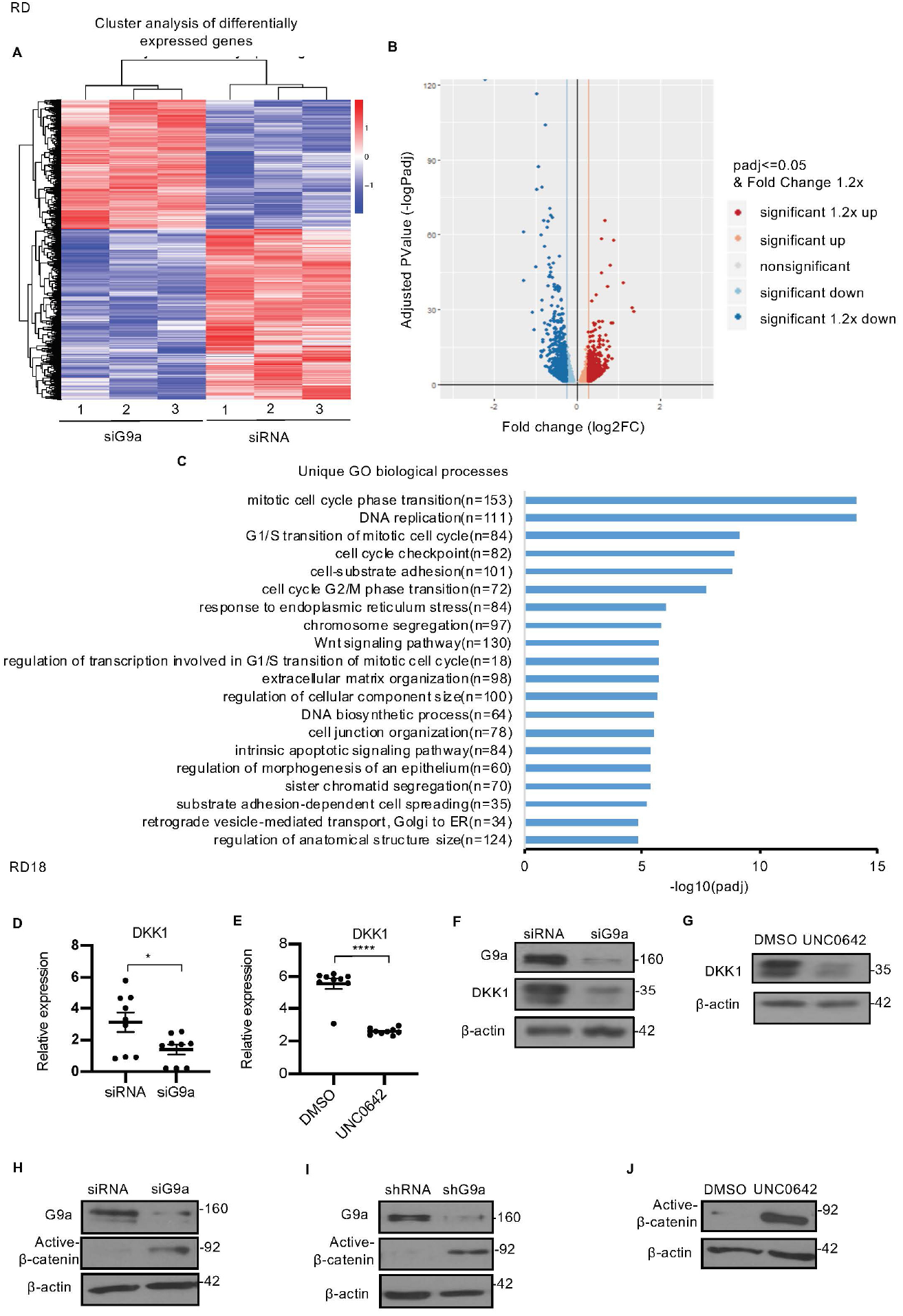
G9a directly regulates DKK1 and Wnt signaling. (**A**) RNA-seq heatmap showing hierarchical clustering of differentially expressed genes. RNA-Seq was performed with control and siG9a RD cells (n=3). Red represents high expression and blue represents low expression. (**B**) Volcano plot showing distribution of differentially expressed genes upon G9a knockdown in RD cells. (**C**) GO enrichment histogram displaying top 20 unique significantly enriched biological processes upon G9a knockdown in RD cells based on *p* adjusted value where n signifies the number of differentially expressed genes concerning the GO term. (**D-E**) qPCR analysis for *DKK1* mRNA in RD18 control and siG9a cells and upon 2.5 μM of UNC0642 treatment (n=3). Values correspond to the average ± SEM. (**F-G**) DKK1 protein was analysed in control and siG9a RD18 cells and in DMSO and 2.5 μM of UNC0642 treated RD18 cells. Representative images of 3 different experiments are shown. (**H-J**) Western blot analysis showed increased active-β-catenin in siG9a cells relative to controls, in stable RD shG9a cells, and upon UNC0642 treatment as indicated. Representative images from 3 different experiments are shown. Statistical significance in **D** and **E** was calculated by unpaired two-tailed *t* test. \**P* ≤ 0.05, \*\*\**P* ≤ 0.001

DKK1 is a member of the Dickkopf family that inhibits canonical Wnt/ß-catenin signaling by binding to and inhibiting the Wnt co-receptor Lrp5/6 (16). Consistent with the transcriptomic data, downregulation of DKK1 mRNA was apparent in siG9a cells compared to control cells by qPCR (Figure 3D). Interestingly UNC0642 treatment also resulted in a decrease in DKK1 mRNA expression (Figure 3E). DKK1 protein levels also decreased in both siG9a and UNC0642 treated cells (Figure 3F and G). Moreover, the reduction in DKK1 levels correlated with increased active-ß-catenin in siG9a cells (Figure 3H), shG9a cells (Figure 3I) and upon UNC0642 treatment (Figure 3J) when compared to their respective controls. These results indicate that loss of G9a leads to downregulation DKK1 with concomitant activation of canonical Wnt signaling and myogenic differentiation.

### G9a regulates DKK1 through Sp1/p300 occupancy

To investigate mechanisms by which G9a activates DKK1 in a methylation dependent manner, we analysed enrichment of the transcription factor Sp1 and the co-activator p300 that have previously been shown to regulate DKK1 expression (24, 25). Sp1 occupancy was detected at the DKK1 promoter (Figure 4A). Intriguingly, both Sp1 and p300 enrichment were decreased upon treatment with UNC0642 compared to control cells. Correspondingly a reduction in H3K9ac, a mark of transcriptional activation was apparent (Figure 4B-D). A decrease in p300 and H3K9ac occupancy was also observed in shG9a cells (Supplemental Figure S4A-B).

**Figure 4.**
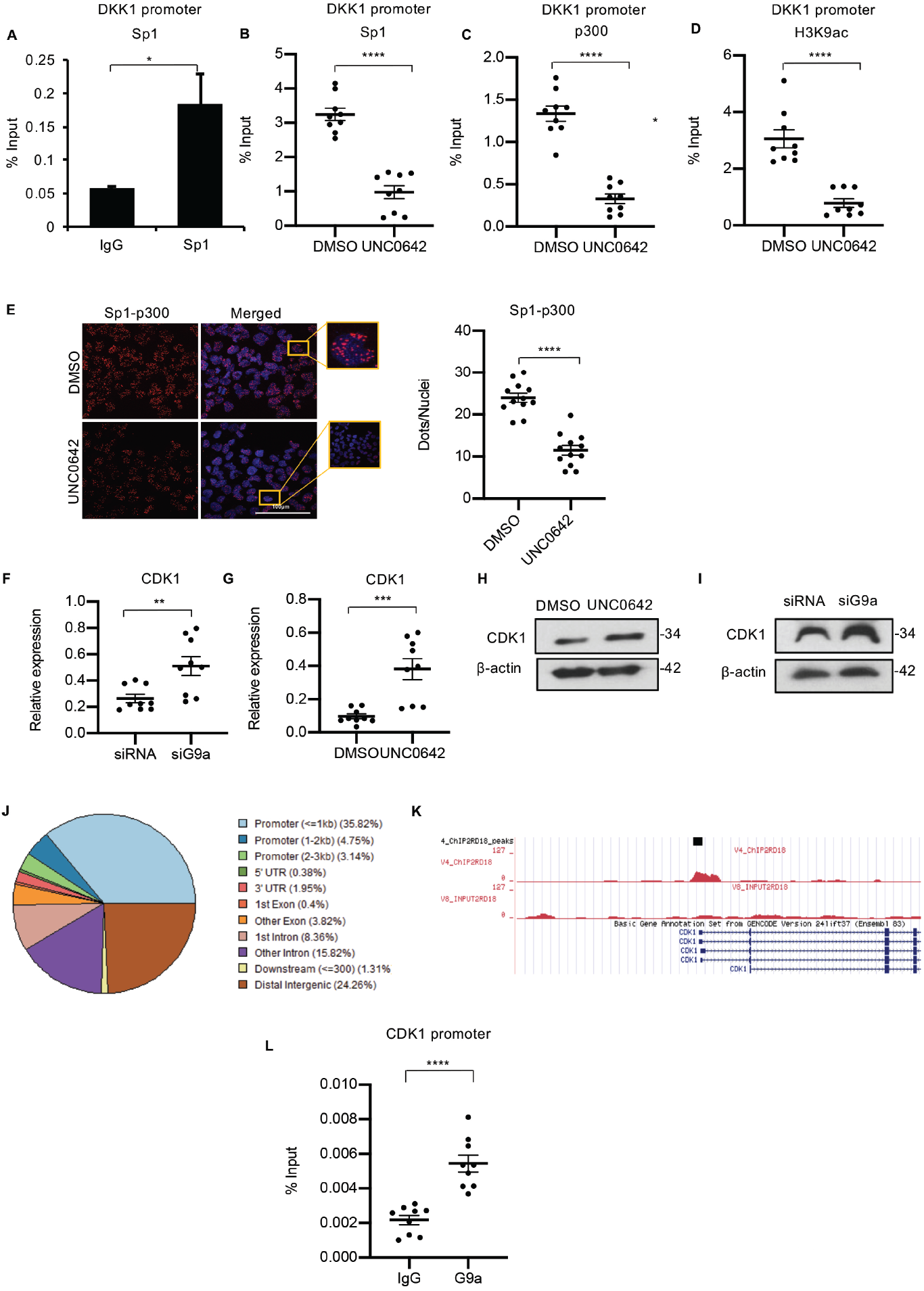
G9a regulates DKK1 expression through Sp1/p300 in a methyltransferase activity dependent manner. **(A)** Sp1 occupancy was analysed by ChIP-PCR at the *DKK1* promoter. IgG was used as a control (n=2). Values correspond to the average ± SD. (**B-D**) Sp1, p300 and H3K9ac enrichment at the *DKK1* promoter was analysed in 2.5 μM of UNC0642 treated cells compared to DMSO controls. The dot plots show reduced enrichment in UNC0642 treated cells (n=3). Values correspond to the average ± SEM. (**E**) PLA was done to examine Sp1 and p300 interaction in control and 2.5 μM of UNC0642 treated cells. Images were captured using confocal microscopy. The dot plot shows the number of dots per nuclei in UNC0642 treated cells compared to control cells (n=3). Each dot represents an interaction. Values correspond to the average ± SEM. (**F-G**) qPCR analysis of *CDK1* mRNA in control and siG9a RD18 cells and in control and 2.5 μM of UNC0642 treated RD18 cells (n=3). Values correspond to the average ± SEM. (**H-I**) CDK1 protein was analysed in in control and siG9a RD18 cells and in DMSO and 2.5 μM of UNC0642 treated RD18 cells. Representative images from 2 independent experiments are shown. (**J**) ChIP-seq analysis in RD18 cells showed G9a occupancy at different regions of the chromatin. (**K**) Snapshot of G9a binding peak at the *CDK1* promoter from the UCSC genome browser. (**L**) G9a occupancy at the *CDK1* promoter was validated by ChIP-PCR (n=3). The dot plot shows enrichment compared to IgG which was used as a control. Values correspond to the average ± SEM. Statistical significance in **A, B, C, D, E, F, G and L** was calculated by unpaired two-tailed *t* test. \**P* ≤ 0.05, \*\**P* ≤ 0.01, \*\*\**P* ≤ 0.001, \*\*\*\**P* ≤ 0.0001.

Sp1 interacts with p300 through its DNA binding domain (26). We therefore examined if Sp1 and p300 interaction was altered by UNC0642 by proximity ligation assay (PLA). The interaction between Sp1 and p300 decreased significantly upon UNC0642 treatment even though that of G9a-Sp1 and G9a-p300 remained unchanged (Figure 4E and Supplementary Figure S4C, D). We then examined mechanisms leading to loss of Sp1 at the DKK1 promoter. Cyclin dependent kinase 1 (CDK1) phosphorylates Sp1 in its DNA binding domain which decreases its DNA binding capacity (27–29). Interestingly, CDK1 was upregulated in siG9a cells in RNA-Seq analysis and qPCR validation confirmed its increase in both siG9a cells and upon UNC0642 treatment at the mRNA and protein level (Figure 4F-I). Similar increase of CDK1 expression upon G9a inhibition was also observed in RD cells (Supplementary Figure 4E-H). To identify if CDK1 is a direct target, ChIP-seq analysis of G9a occupancy was performed which revealed its binding mostly at gene promoters (Figure 4J) and its enrichment was apparent at the *CDK1* promoter (Figure 4K). To validate these results, ChIP-PCR was done in RD18 cells and a significant enrichment was seen indicating that G9a directly binds to the *CDK1* promoter (Figure 4L). These results indicate that G9a represses *CDK1* expression in a methyltransferase activity-dependent manner. Upregulation of *CDK1* expression upon G9a inhibition may result in increased Sp1 phosphorylation and its decreased DNA binding resulting in reduced Sp1-p300 occupancy and consequently downregulation of *DKK1* expression.

### G9a inhibits differentiation and promotes proliferation through DKK1-mediated antagonism of Wnt signaling

Canonical Wnt signaling induces myogenic differentiation and decreases proliferation (30, 31). As DKK1 is a well characterized inhibitor of canonical Wnt signaling, we examined if the effect of G9a is mediated by DKK1. Correlating with high endogenous G9a expression, DKK1 was also overexpressed in all three lines compared to HSMM at both mRNA and protein level (Figure 5A and B). Moreover, analogous to G9a knockdown, DKK1 knockdown in RD18 cells resulted in a significant decrease in BrdU^+^ cells compared to control cells (Figure 5C). A corresponding increase in differentiation was also apparent by elevated MHC levels and myogenin expression (Figure 5D). To determine whether DKK1 mediates the effects of G9a, we performed rescue experiments. Recombinant DKK1 (rDKK1) was added to siG9a cells for 24 hr that resulted in the reduction of active β-catenin seen in siG9a cells (Figure 4E). Interestingly, in presence of rDKK1, the increase in MHC^+^ cells and myogenin expression in siG9a cells was reversed to control levels (Figure 5E). Similarly, the decrease in BrdU^+^ cells upon G9a knockdown were restored to levels comparable to control. To further validate that G9a mediates its effects on canonical Wnt signaling, we used another Wnt antagonist, a porcupine inhibitor, LGK974. Similar to rDKK1, LGK974 reversed the effects of G9a knockdown on proliferation, differentiation and active β-catenin levels (Figure 5F) indicating that G9a mediates the differentiation block by activating *DKK1* expression that in turn suppresses canonical Wnt signaling.

**Figure 5.**
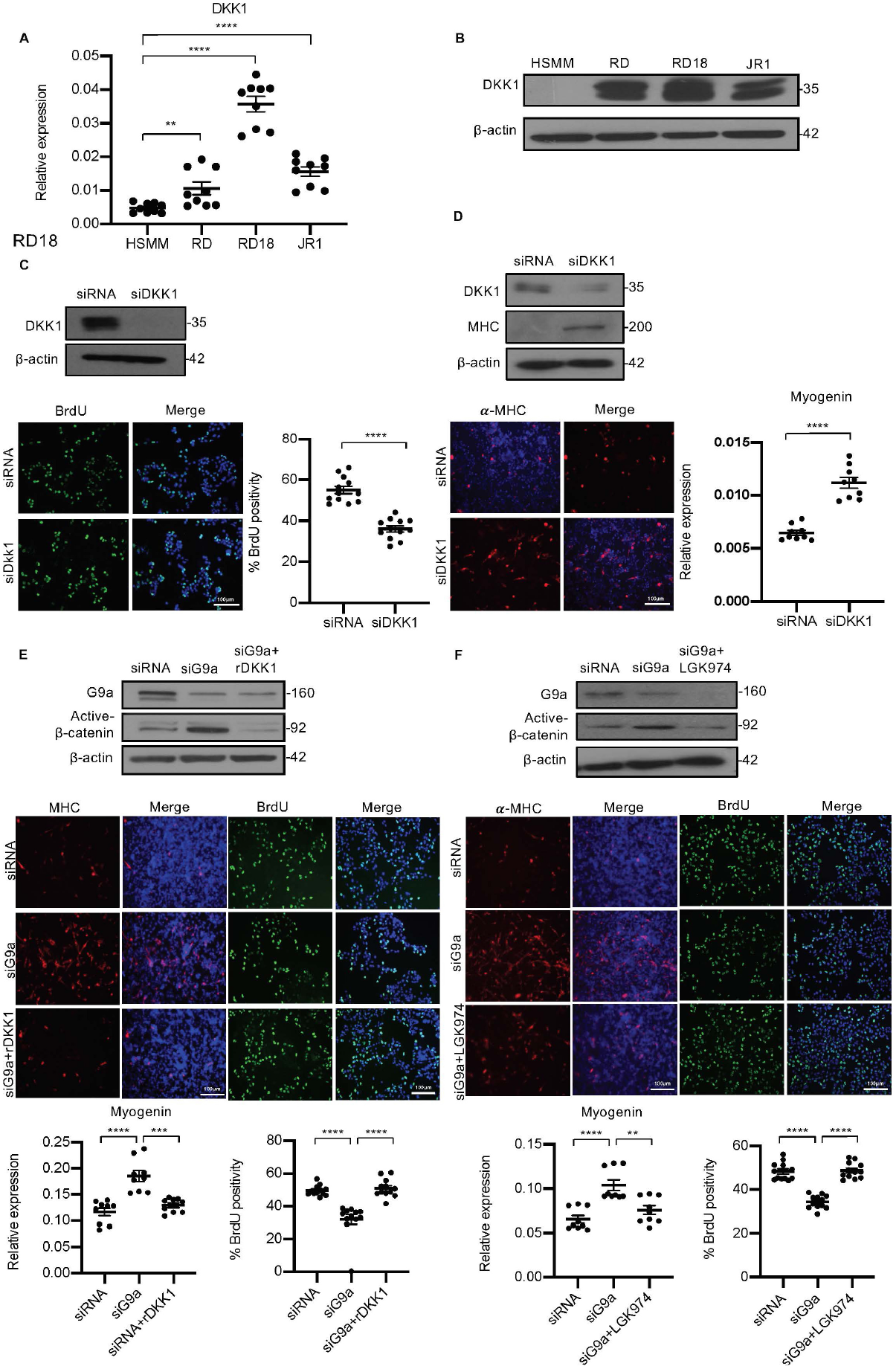
DKK1 is a downstream effector of G9a function. (**A**) *DKK1* mRNA was examined by qPCR (n=3) in HSMM, RD, RD18 and JR1 cells. Values correspond to the average ± SEM. (**B**) DKK1 protein levels were analysed by western blotting. A representative image from 3 different experiments is shown. DKK1 knockdown was analysed in control and siDKK1 cells by westen blot. Proliferation was analysed in RD18 control and siDKK1 cells (n=3) with anti-BrdU antibody. The dot plot shows the percentage of BrdU^+^ cells. Values correspond to the average ± SEM. (**D**) Differentiation was analysed in control and siDKK1 cells that were cultured for five days in DM. Cells were analysed by western blot and immunofluorescence and using anti-MHC antibody as indicated. A representative image of 3 different experiments is shown. Myogenin expression was analysed by qPCR (n=3) at day 2 of differentiation. Values correspond to the average ± SEM. (**E**) Control, siG9a cells and siG9a cells treated with rDKK1 for 24 hr and tested for active-β-catenin levels. Differentiation and proliferation was analyzed (lower panels) by MHC^+^ cells and BrdU^+^ cells as indicated. Representative images of 3 different experiments are shown. Myogenin expression was analysed by qPCR (n=3) and the percentage of BrdU^+^ cells is shown in the dot plots. Values correspond to the average ± SEM. (**F**) Western blot showing active-β-catenin levels in control, siG9a cells and siG9a cells treated with LGK974 for 24 hr. A representative image of 3 different experiments are shown. MHC^+^ and BrdU^+^ cells were analysed. A representative image of 3 different experiments is shown. Myogenin expression in control, siG9a and siG9a RD18 cells treated with LGK974 was analysed by qPCR (n=3). Values correspond to the average ± SEM. Statistical significance in **A, C, D, E, and F** was calculated by unpaired two-tailed *t* test. \*\**P* ≤ 0.01, \*\*\**P* ≤ 0.001, \*\*\*\**P* ≤ 0.0001.

In order to examine the effect of G9a in regulating DKK1 and Wnt signaling *in vivo*, we injected RD cells in BALB/c nude mice. Once the tumors were palpable, mice were injected intraperitoneally every two days with UNC0642 or with control vehicle. Treatment with UNC0642 resulted in reduced tumor growth compared to the control group without any significant changes in body weight (Figure 6A). By immunohistochemical analysis (Figure 6B) we confirmed a decrease in H3K9me2 in tumors from mice treated with UNC0642 indicating efficacy of the drug *in vivo*. The proliferation marker Ki67 was decreased, whereas MHC^+^ cells were increased in tumors from mice treated with UNC0642. Moreover, DKK1 was decreased upon UNC0642 treatment, and correspondingly active-ß-catenin levels were elevated (Figure 6B).

**Figure 6.**
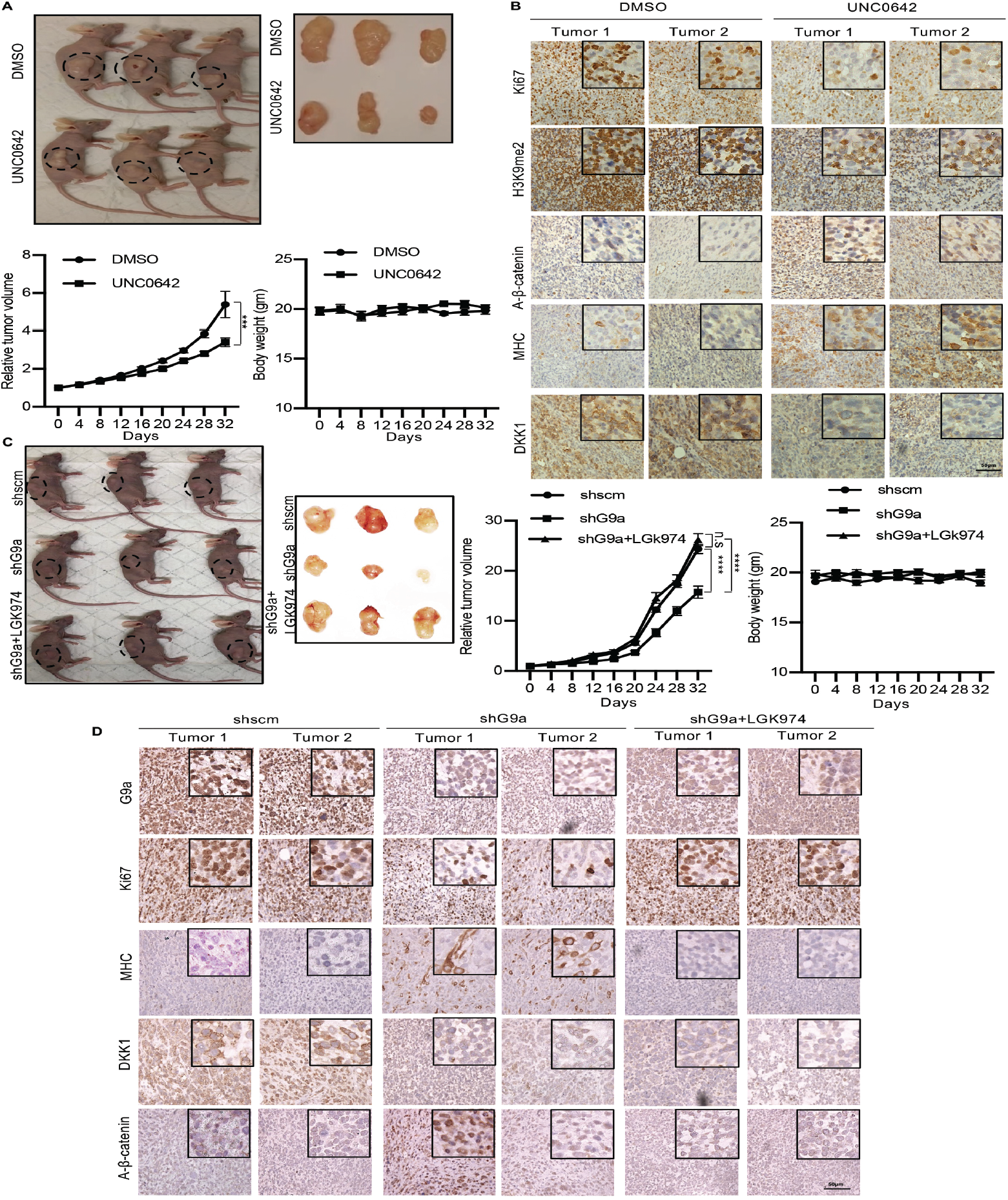
G9a regulates tumor growth by regulation of DKK1 and Wnt signaling. (**A**) Nude mice were injected with RD cells. Once tumours were palpable, mice were treated with DMSO (n=10) or UNC0642 (n=10). Representative images of three mice in each group (left panel), and resected tumors (right panel) are shown. The relative tumor volume in UNC0642 treated group showed a significant decrease compared to controls although the body weight of mice did not show any significant change. Statistical significance was calculated using repeated-measure two-way ANOVA where ***P ≤ 0.001. Values correspond to the average ± SEM. (**B**) Tumors from 2 control and 2 UNC0642 treated mice were analysed by IHC using anti-Ki67, anti-H3K9me2, anti-active-ß-catenin anti-MHC and anti-DKK1 antibodies. Scale bar: 50μm. Inset shows zoomed in images. (**C-D**) Mice were injected with shRNA cells (shscm) (n=10) or shG9a cells (n=20). Once tumors were palpable, half of shG9a injected mice were treated with vehicle and the rest with LGK974. (**C**) Representative images of mice (left panel) injected with shscm control cells, shG9a cells and shG9a cells + LGK974 are shown. Representative images of the tumors (right panel) isolated from the three cohorts are shown. The relative tumor volume and the body weight of mice in each group was determined. Statistical significance was calculated using repeated-measure two-way ANOVA where ****P ≤ 0.0001. Values correspond to the average ± SEM. Tumors from two different mice in each group were analyzed by IHC for Ki67, H3K9me2, DKK1, MHC and active-ß-catenin staining as described above. Scale bar: 50μm. Inset shows zoomed in images.

To verify that the effects of G9a are mediated via an impact on Wnt signaling *in vivo*, we next injected control shRNA and shG9a cells. Once the tumors were palpable, mice injected with shG9a cells were treated with LGK974 or treated with vehicle alone every alternate day. The shRNA control group was also injected with the vehicle (Figure 6C). Tumor volume in mice injected with shG9a cells was reduced compared to the control group. However, mice injected with shG9a cells and treated with LGK974 showed tumor volumes comparable to the control group. Body weight of mice did not show any significant changes over the course of this treatment. We analysed two tumors from each cohort by IHC. As expected, G9a expression was decreased in mice injected with shG9a cells that correlated with decrease in Ki67^+^ cells and increase in MHC^+^ cells compared to tumors of the control group. This alteration was however not seen in the tumors from mice injected with shG9a cells that underwent LGK974 treatment. Consistently, the reduction of DKK1 and increase in active-ß-catenin in tumors from shG9a injected mice was reversed in upon LGK974 treatment (Figure 6D). Taken together, these studies indicate that inhibition of G9a expression or activity decreased tumor progression by regulating Wnt signaling.

## Discussion

In this study, we uncovered a G9a-dependent epigenetic node that results in repression of Wnt signaling in ERMS. We propose that by activating DKK1 expression, G9a maintains Wnt signaling in a repressed state, and thus prevents the transition of myoblasts to a differentiated state. These findings underscore specific epigenetic mechanisms to reactivate Wnt signaling and induce differentiation in ERMS.

There has been a resurgence of interest in differentiation therapy as a viable treatment option for solid tumors (32). The Wnt, Notch and Hedgehog pathways play a pivotal role major role in balancing proliferation, self-renewal and differentiation during embryonic myogenesis. Not surprisingly, deregulation of these developmental pathways has been reported in ERMS (12, 33, 34) and consequently, drugs targeting each of these pathways are being tested. However, current inhibitors do not selectively target specific pathways and have either unacceptable toxicity, or do not show marked clinical improvement. Several studies have demonstrated suppression of Wnt signaling in RMS cells, which mostly do not stain positively for nuclear β-catenin. Also, no mutations in the β catenin gene have been reported (35, 36). The suppression of Wnt signaling is a critical contributor to the differentiation block in ERMS, and induction of the pathway leads to cell cycle exit and differentiation (12, 14). GSK3ß inhibitors are the most prominent Wnt signaling activators and showed promising effects in inducing differentiation in a zebrafish model of ERMS (14). GSK3ß is a constitutive serine/threonine protein kinase that inhibits canonical Wnt signaling by phosphorylating ß-catenin and triggering its degradation in the cytoplasm (37). However, GSK3ß is involved in many pathways with more predicted substrates than any known kinase. Thus, its inhibition is not a specific strategy to induce Wnt signaling. This is also emphasized by slow progress of existing GSK3ß inhibitors toward clinical translation (38). Tideglusib, an irreversible GSK3ß inhibitor, was recently tested against rhabdomyosarcoma PDX models, where the highest safe dosage failed to both induce myogenic differentiation as well as affect cancer progression in PDX models (39). Consequently, development of alternative molecularly targeted therapies that induce Wnt signaling is critical goal in this disease.

Our data demonstrates an epigenetic mechanism to activate Wnt signaling and overcome the differentiation block in ERMS. Interestingly G9a activates DKK1 in a methylation-dependent manner resulting in the suppression of Wnt signaling in ERMS. While methylation-dependent silencing or repression by G9a has been described, only a few studies have demonstrated methylation-dependent activation of gene expression. Consistent with previous studies, we found Sp1 binding at the *DKK1* promoter and inhibition of G9a led to its decreased occupancy. Sp1 transcriptional activity is regulated through a variety of post-translational modifications (27). CDK1/cyclin B1 phosphorylates Sp1 at Threonine (Thr) 739 at the Sp1 DNA binding site (29) and decreases its DNA binding. Since *CDK1* is repressed by G9a in a methyltransferase-dependent manner, its elevated levels upon UNC0642 treatment may result in reduced Sp1 binding. Phosphorylation of Sp1 at the DNA binding domain might also result in the dissociation of p300 from Sp1, as Sp1 interacts with p300 through its DNA binding domain (26). Consequently, the activation mark H3K9ac also decreased at the *DKK1* promoter. Thus, our data establish a novel mechanism in which G9a methyltransferase activity is required for Sp1-p300 association for activation of *DKK1* expression. However, there may be additional mechanisms by which G9a regulates DKK1 expression.

Pan-HDAC and pan-DNMT inhibitors have been explored in ERMS (9, 40). HDAC inhibitors often exert effects independent of their epigenetic roles (9) and application of DNMT inhibitors (41) are restricted due to their toxicity in healthy cells. EZH2 inhibitors are the only other specific epigenetic inhibitors that demonstrate a strong phenotype in ERMS (8). In this study, a drug screen of 15 methyltransferase inhibitors in two different ERMS cell lines showed that small molecule inhibitors targeting G9a activity are very effective. Thus, our data support targeting G9a as a therapeutic approach in ERMS, particularly since its deletion does not impact development of muscle (42). We have recently shown that G9a is deregulated in ARMS as well (20). Together with the herein described role of G9a in ERMS, these observations clearly suggest the importance of differentiation in these therapies and imply common features between these two subtypes of RMS.

## Materials and Methods

### Cell culture and drug sensitivity assays

RD, RD18 and JR1 ERMS cell lines were a kind gift from Peter Houghton (Nationwide Children’s Hospital, Ohio, USA) and Rosella Rota (Bambino Gesu Children’s Hospital, Rome, Italy). RD18 and JR1 were cultured in RPMI 1640 with L-Glutamine (Thermo Fisher Scientific, Waltham, MA, USA) and 10% FBS (Hyclone, Logan UT, USA), whereas RD cells were cultured in Dulbecco’s Modified Eagle Medium (DMEM) (Sigma, St Louis, MO, USA) with 10% FBS (Hyclone, Logan UT, USA). Primary human skeletal muscle myoblasts (HSMMs) were purchased from Lonza Inc. (Basel, Switzerland) and cultured in growth medium (SkGM-2 BulletKit). For transient knockdown, cells were transfected with 50nM of human G9a-specific siRNA or human DKK1-specific siRNA (ON-TARGETplus siRNA SMARTpool, Dharmacon, Lafayette, CO, USA) containing a pool of three to five 19–25 nucleotide siRNAs. Control cells were transfected with 50nM scrambled siRNA (ON-TARGETplus, non-targeting pool, Dharmacon) using Lipofectamine RNAiMax (Thermo Fisher scientific). Cells were analysed 48 hr post-transfection for all assays. Knockdown efficiency was monitored by western blot. For generating stable knockdown cell lines, RD cells at 40-50% confluency were transduced with shRNA control lentivirus particles (Santa Cruz Biotechnology Inc.), or shG9a lentivirus particles (Santa Cruz Biotechnology Inc.) and 2μl polybrene (8mg/ml) (Sigma-Aldrich) in DMEM basal medium. 6 hr post-transduction, cell supernatants were replaced with DMEM medium (10% FBS) for 24 hr. Transduced cells were selected with 1μg/ml puromycin (Sigma-Aldrich) for four days. For rescue experiments, siG9a cells were treated with 100ng/ml of rDKK1 (R&D Systems) or 200nM of porcupine inhibitor LGK974 (Selleck Chemicals, Houston, USA). Both rDKK1 and LGK974 were added to the media 24 hr after transfection. For drug screening, RD and JR1 cells were treated with methyltransferase inhibitors at 3, 1, and 0.3 μM. The number of viable cells at day 8 was examined by MTS assay, and calculated as the ratio over control cells treated with an equivalent dilution of DMSO. The data are presented as a heatmap where red indicates viability ratio>control; white=control; and blue is less than control. The experiment was conducted in triplicate and (+)-JQ1 was used as a positive control.

### Proliferation and differentiation assays

Proliferative capacity of cells was analysed using 5-bromo-2’-deoxy-uridine (BrdU) labelling (Roche, Basel, Switzerland). Cells seeded on coverslips were pulsed with 10 μM BrdU for 60 min at 37°C. Cells were fixed with 70% ethanol at −20°C for 20 min and incubated with anti-BrdU antibody (1:100) for 60 min followed by anti-mouse Ig-fluorescein antibody (1:200) for 60 min. After mounting onto slides with DAPI (Vectashield, Vector Laboratories, CA, USA), images were captured using fluorescence microscope BX53 (Olympus Corporation, Shinjuku, Tokyo, Japan). For differentiation assays, RD, RD18 and JR1 cells were cultured for 2-5 days in DMEM supplemented with 2% horse serum (Gibco, Carlsbad, CA, USA) at 90-95% confluency. Differentiation was assessed by MHC staining. Cells were fixed with 4% paraformaldehyde for 20 min at room temperature (RT). Cells were blocked and permeabilised using 10% horse serum and 0.1% Triton X containing PBS. Cells were then incubated with anti-Myosin Heavy Chain (MHC) primary antibody (R&D Systems, Minneapolis, MN, USA) (1:400, 1 hr at RT) followed by 1 hr of 1:250 secondary goat anti-Mouse IgG (H+L) Highly Cross-Adsorbed Secondary Antibody, Alexa Fluor 568 (Thermo Fisher scientific). Coverslips were mounted with DAPI (Vectashield, Vector Laboratories, CA, USA) and imaged using upright fluorescence microscope BX53 (Olympus Corporation).

### Western blot analysis

Cells were lysed using RIPA or SDS lysis buffer supplemented with protease inhibitors (Complete Mini, Sigma-Aldrich). The following primary antibodies were used: anti-G9a (#3306S, 1:300, Cell Signaling) anti-MHC (#sc-32732, 1:300, Santa Cruz Biotechnology), anti-Myogenin (#sc-12732, 1:250, Santa Cruz Biotechnology), anti-DKK1(#sc374574, 1:300, Santa Cruz Biotechnology), anti-active-ß-catenin (#05-665, 1:500, Merck Millipore), anti-H3K9me2 (#9753S, 1:1000, Cell Signaling), anti-ß-actin (#A2228, 1:10,000; Sigma-Aldrich), anti-H3 (#ab1791, 1:10,000; Abcam). The primary antibody against CDK1 was home grown and a kind gift from Philip Kaldis (A* Star, Institute of Molecular and Cellular Biology). Appropriate secondary antibodies (IgG-Fc Specific-Peroxidase) of mouse or rabbit origin (Sigma Aldrich) were used.

### Proximity Ligation Assay

Proximity ligation assay (PLA) was performed using the Duolink in situ-fluorescence (Sigma DUO92101). For G9a and Sp1 interaction, anti-G9a (Cell Signaling, 1:50) and anti-Sp1(1:50 Santa Cruz) antibodies were used. For Sp1 and p300 interaction studies, Sp1 (Millipore, 1:100) and p300 (Abcam, 1:1000) antibodies were used. Images were captured under FluoView FV1000 confocal fluorescence microscope (Olympus) at 60X (oil). For quantifying PLA signals, particle analysis was performed using Fiji/ImageJ software, and pixel area size of 2-50 was assigned for calculating the total number of PLA signals per field. PLA signals as dots per nuclei were calculated for at least 3 microscopic fields.

### Transcriptome analysis and Quantitative real-time polymerase chain reaction (qPCR)

For RNA sequencing analysis, RNA was isolated from control and siG9a cells in triplicate using Trizol. RNA was sequenced using Illumina high-throughput sequencing platform. CASAVA base recognition was used to convert raw data file to Sequence Reads and stored in FASTQ(fq) format. Raw reads were then further filtered in order to achieve clean reads using the following filtering conditions: reads without adaptors, reads containing number of base that cannot be determined below 10% and at least 50% bases of the reads having Qscore denoting Quality value <=5. For mapping of the reads STAR software was used to align the reads against hg19 Homo sapiens reference genome. 1M base was used as the sliding window for distribution of the mapped reads. For analysis of the differentially expressed genes, Gene Ontology analysis was done using cluster Profiler (43) software for GO terms with corrected *p* value less than 0.05.

For qPCR analysis, total RNA was extracted using Trizol (Thermo Fisher Scientific) and quantified using Nanodrop. Messenger RNA (mRNA) was converted to a single-stranded complementary DNA (cDNA) using iScript cDNA Synthesis Kit (Bio-Rad). qPCR was performed using Lightcycler 480 SYBR Green 1 Master Kit (Roche). PCR amplification was performed as follows: 95 °C 5 min, followed by 95 °C for 10s, annealing at 60 °C for 10s, followed by 45 cycles at 72 °C for 10s. Light Cycler 480 software (version 1.3.0.0705) was used for analysis. CT values of samples were normalized to internal control GAPDH to obtain delta CT (ΔCT). Relative expression was calculated by 2−ΔCT equation. qPCR was done using reaction triplicate and at least two independent biological replicates were done for each analysis. Primer sequences for G9a are: 5’- TGGGCCATGCCACAAAGTC-3’ and 5’- CAGATGGAGGTGATTTTCCCG-3’, for Myogenin are 5’-GCCTCCTGCAGTCCAGAGT-3’ and 5’- AGTGCAGGTTGTGGGCATCT-3’, for DKK1 are 5’-CGGGAATTACTGCAAAAATGGA-3’ and 5’- GCACAGTCTGATGACCGGAGA-3’ and CDK1 are 5’-TTTTCAGAGCTTTGGGCACT-3’ and 5’- CCATTTTGCCAGAAATTCGT-3’.

### Chromatin immuno-precipitation (ChIP)

Chromatin immunoprecipitation-sequencing (ChIP-Seq) was done using 20 million RD18 cells and anti-G9a antibody (Abcam, Cambridge, MA, USA) as described (20). Sequencing reads were mapped against human reference genome hg19. High quality mapped reads (MAPQ>=10) were retained and potential duplicates were removed using SAMtools. G9a binding sites were predicted from the libraries using MACS2. Read density was computed in the format of bigwig using MEDIPS with 50bp window width. The prediction revealed 48,999 binding sites overlapping with promoters, gene body and intergenic regions. We used GENCODE v19 to define promoters (+/− 2.5kb from TSS) and gene body. Mid-point predicted binding sites were used in this analysis. Of the 48,999 binding sites 49% (n=24,176) localized at promoters, 29% (n=14,252) at gene bodies and 21% (n=10,571) at inter-genic regions. To demonstrate the read density around annotated TSS, we identified the promoters of the TSS overlapping with predicted G9a binding sites. The TSS were then extended +/− 20kb and the binding signal was computed in each window of size 100bp. Average read density for each window was computed using bigWigAverageOverBed. GENECODEv19 was used to classify promoter-bound G9a binding. Peaks were annotated using ChIPseeker (44) and ChIPpeakAnno (45) against genes model of UCSC, hg19, knownGene (TxDb.Hsapiens.UCSC.hg19.knownGene). Differential Expression count matrix were analysed using R. Genes with an adjusted p-value less than 0.05 were labelled as significantly differentially expressed. From that list, genes with an absolute fold change >= 1.2 were selected for further analysis. GO analysis of gene subsets were performed using Metascape (46). The ChIP-Seq data are compliant with MIAME guidelines and have been deposited in the NCBI GEO database.

ChIP-PCR was done as previously described (20). Relative enrichment was calculated using 2−ΔCT equation. The following antibodies were used for ChIP assays: ChIP-grade anti-G9a (Abcam), anti-H3K9ac (Abcam), Sp1 (Rabbit Millipore) and p300 (Abcam). Primers used for ChIP at the *DKK1* promoter were: Forward: 5’-CCGGATAATTCAACCCTTACTGCC-3’ and Reverse: 5’- GGAGCATTCCGGCCCCTTGGGAG-3’; and for ChIP at the *CDK1* promoter were: Forward: 5’- CCACACTGGGCTGGCTTTAG-3’ and Reverse: ‘5-AGTTCCTGAGAACAGCCGAC-3’.

### Mouse xenograft experiments

6-week-old C.Cg/AnNTac-Foxn1^nu^NE9 female BALB/c nude mice (InVivos, Singapore) were injected subcutaneously in the right flank with control RD cells (10 × 10^6^). Once tumors were palpable, one group (n=10/group) was injected intraperitoneally with vehicle [5% DMSO in PBS], and the other with UNC0642 (5mg/kg body weight in 5% DMSO) every alternate day. Tumor diameter and volume was calculated as described (20). Resected tumors were fixed and paraffin sections were immunostained with various antibodies. To determine the role of Wnt signaling, one group of mice were injected with

RD shcontrol cells (n=10/group) and two groups with RD shG9a cells. Once tumors were palpable, the control group and one shG9a group were injected intraperitoneally with control vehicle [2% DMSO in corn oil], and one group of shG9a mice was injected with LGK974 (5mg/kg body weight in 2% DMSO). Tumour growth and body weight were recorded as described (20). All animal procedures were approved by the Institutional Animal Care and Use Committee.

### Immunohistochemistry (IHC)

Paraffin sections from 16 primary ERMS archival tumour specimens and three normal muscle from National University Hospital (NUH) and KK Women’s and Children Hospital in Singapore were analysed by IHC using anti-G9a antibody (1:50 dilution, Cell Signaling) as described (20). Negative controls were performed using secondary antibody only. Images were captured with Olympus BX43 microscope (Ina-shi, Nagano, Japan). Approval was obtained from the ethics committee (IRB) at NUS. For IHC on mouse xenografts, sections were incubated with anti-G9a (1:200, Abcam), anti-H3K9me2 (Abcam), anti-Ki67 (1:100, Santa Cruz Biotechnology), anti-active-ß-catenin (1:300; Merck Millipore), anti-MHC (#M4276 1:200 Sigma-Aldrich), anti-DKK1 (#ab61034 Abcam) antibodies followed by biotinylated goat anti-rabbit/anti-mouse IgG (H+L) secondary antibody (Vector Laboratories). Sections were washed and incubated with Vectastain Avidin–Biotin Complex (Vector Laboratories) for 20 min at 37°C.

#### Statistical analysis

For statistical analysis, two-tailed non-parametric unpaired *t* test was used to evaluate significance with the use of GraphPad prism 9.0 software. Each experiment had three biological replicates. Standard error of mean (SEM) was calculated for all data sets and a *p* value less than 0.05 was considered statistically significant. \**P* ≤ 0.05, \*\**P* ≤ 0.01, \*\*\**P* ≤ 0.001, \*\*\*\**P* ≤ 0.0001. For *in vivo* experiments repeat-measure two-way ANOVA was used to calculate the statistical significance between different groups.

## Data availability

ChIP-Seq data has been deposited in GEO under the accession number GSE125960. Private token for referees: qbgtkgganjmlrwd. RNA-Seq data been deposited in GEO under the accession number GSE142975. Private token for referees: unylciiwfvorjgd.

## Authors’ Contributions

Conception and design: A Pal, R Taneja

Development of methodology: A Pal, JY Leung, G Ang, VK Rao, HJ Lim, L Pignata

Acquisition of data (provided animals, acquired and managed patients, provided facilities, etc.): VLK Min, KTE Chang

Analysis and interpretation of data (e.g., statistical analysis, biostatistics, computational analysis): VLK Min, M Hebrard, E Guccione, R Taneja

Writing, review, and/or revision of the manuscript: A Pal, L Pignata, E Guccione and R Taneja Administrative, technical, or material support (i.e., reporting or organizing data, constructing databases): A Pal, VK Rao, M Hebrard

Study supervision: R Taneja

## Acknowledgements

We thank Peter Houghton and Rosella Rota for ERMS cell lines; Ooi Wen Fong, Genome Institute of Singapore for bioinformatics analysis; Philip Kaldis for CDK1 antibody, and David Virshup Duke-NUS for helpful discussions. This work was supported by a National Medical Research Council grant (NMRC/OFIRG/0073/2018) to R.T. A.P is supported by the President’s Graduate Scholarship at the National University of Singapore.

---

**Supplemental Figure 1.**
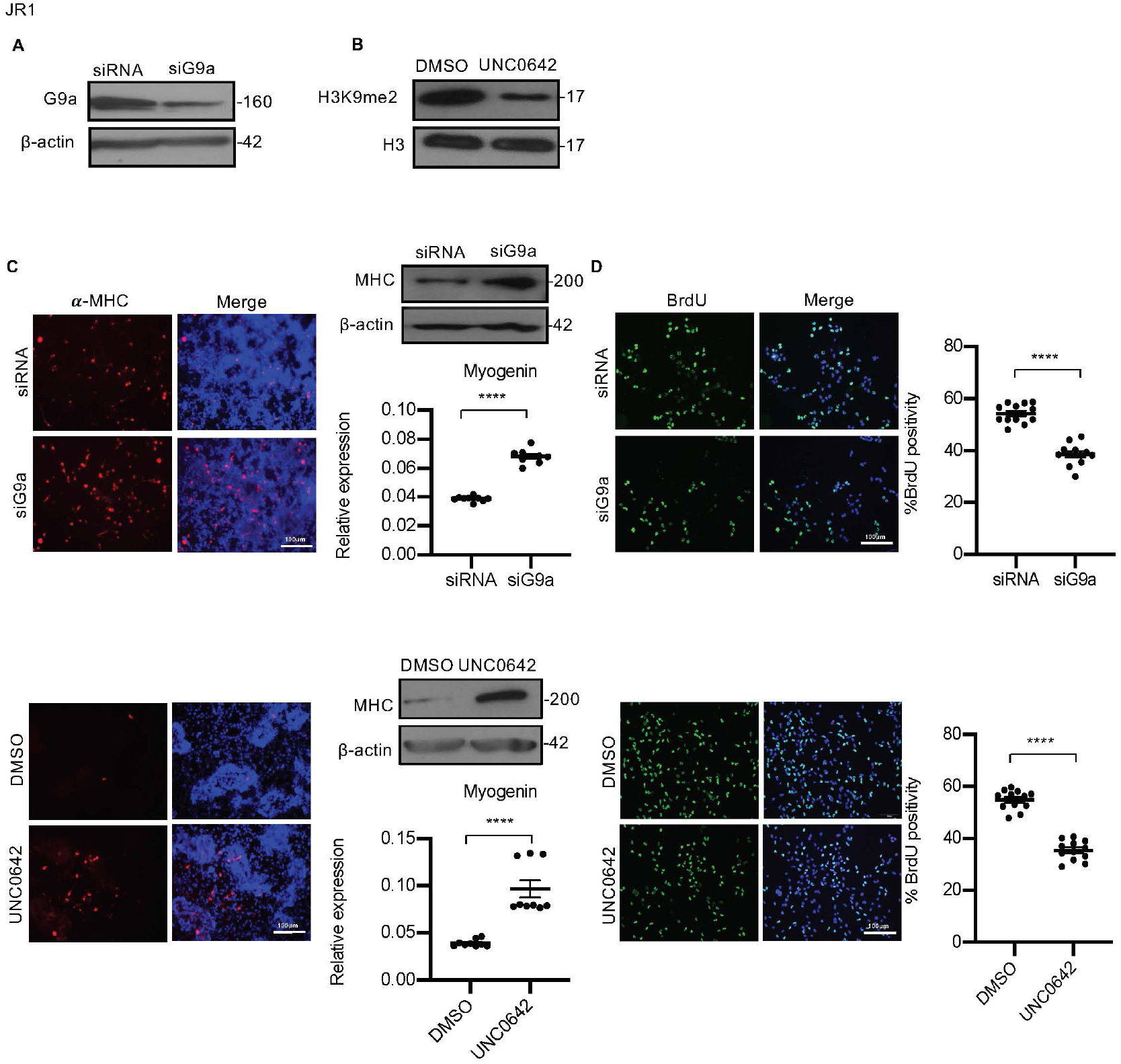
Loss of G9a expression or activity in JR1 cells increases differentiation and reduces proliferation. **(A)** Western blot showing G9a knockdown efficiency in siG9a cells. (**B**) Inhibition of G9a activity by UNC0642 decreased H3K9me2 protein level after 48 hr of treatment as shown by western blot. (**C**) G9a knockdown and UNC0642 treatment resulted in increased MHC^+^ cells as seen by immunofluorescence assay and western blot after five days in DM. Data is representative of 3 independent experiments. G9a knockdown and treatment with UNC0642 also increased myogenin expression as seen by qPCR analysis (n=3). Values correspond to the average ± SEM. (**D**) G9a knockdown and UNC0642 treatment reduced BrdU^+^ cells as seen by immunofluorescence assays. Data is representative of 3 independent experiments. The dot plots show the percentage of BrdU^+^ cells in siG9a cells and UNC0642 treated cells relative to controls. Values correspond to the average ± SEM. Statistical significance in **C** and **D** was calculated by unpaired two-tailed *t* test. \*\*\*\**P* ≤ 0.0001.

**Supplemental Figure 2.**
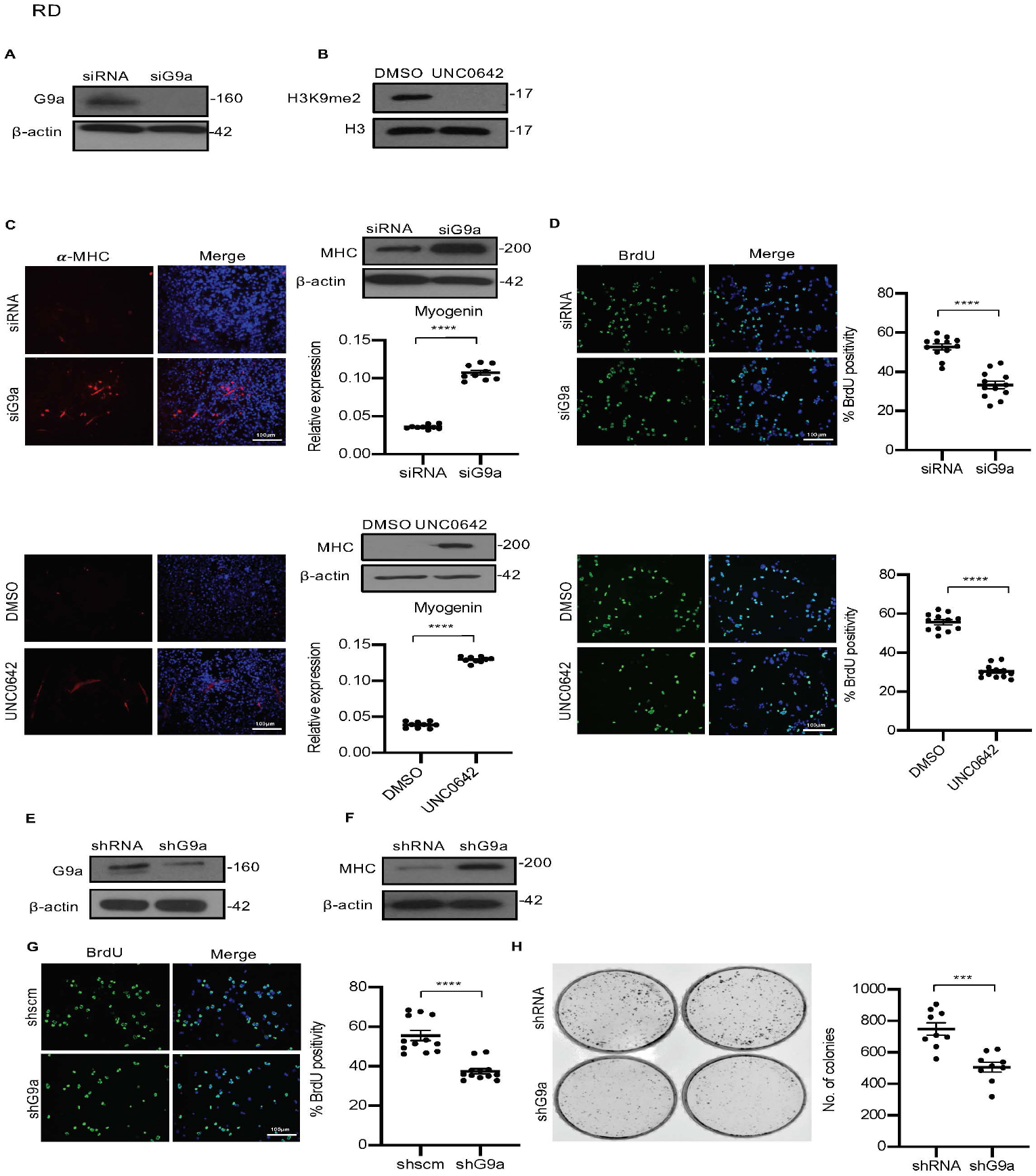
Loss of G9a expression or activity in RD cells increases differentiation and reduces proliferation. **(A)** Western blot showing G9a knockdown efficiency. **(B)** Inhibition of G9a activity by UNC0642 decreased H3K9me2 protein level after 48 hr of treatment as shown by western blot. **(C)** G9a knockdown and treatment with UNC0642 increased MHC^+^ cells as seen by immunofluorescence assay and western blot after five days in DM. The image is representative of 2 independent experiments. G9a knockdown and treatment with UNC0642 also increased myogenin expression as seen by qPCR analysis (n=3). Values correspond to the average ± SEM. (**D)** G9a knockdown and UNC0642 treatment reduced BrdU^+^ cells 48 hr in growth media as seen by immunofluorescence assays. The dot plots show the percentage of BrdU^+^ cells relative to controls (n=3). Values correspond to the average ± SEM. **(E)** Lentiviral mediated stable G9a knockdown (shG9a) efficiency in RD cells is shown by western blot. **(F)** shG9a cells showed increased MHC levels after five days in DM by western blot. Data is representative of 2 independent experiments. **(G)** shG9a cells showed reduced BrdU^+^ cells compared to control. Data is representative of 3 independent experiments. The percentage of BrdU^+^ cells is shown. Values correspond to the average ± SEM. **(H)** Colony formation was analysed in control and shG9a cells. The number of colonies was quantified which showed reduced numbers in shG9a cells. (n=3). Values correspond to the average ± SEM. Statistical significance in **C**, **D, G and H** was calculated by unpaired two-tailed *t* test. \*\*\**P* ≤ 0.001 and \*\*\*\**P* ≤ 0.0001.

**Supplemental Figure 3:**
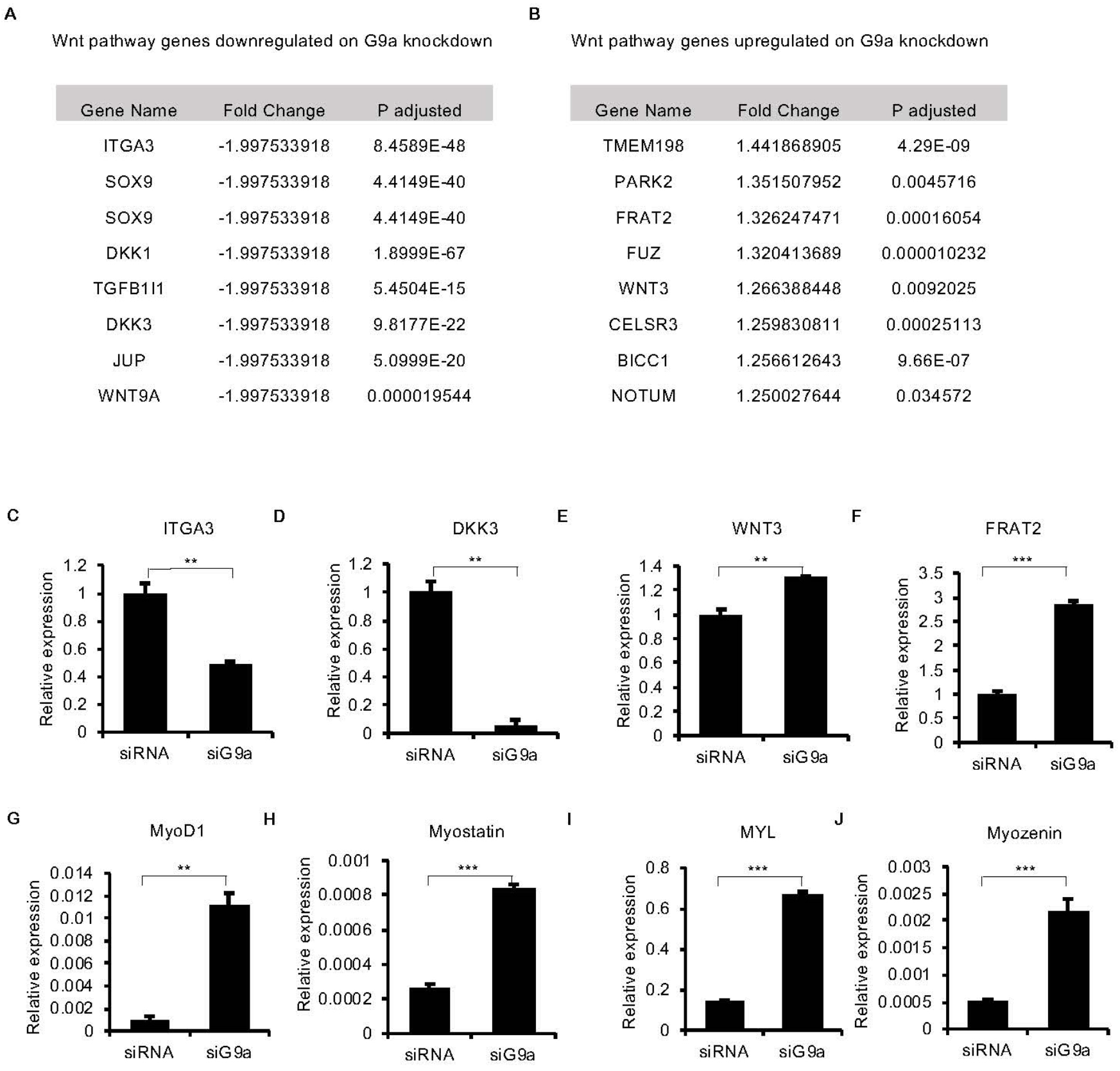
Validation of RNA sequencing analysis. (**A-B**) List of top significantly altered Wnt pathway genes identified in RNA sequencing analysis that are downregulated, or upregulated, upon G9a knockdown. **(C-F)** qPCR validation of 2 downregulated genes ITGA3 and DKK3 and 2 upregulated genes FRAT2 and WNT3 in RD cells. Data shown is representative of 2 independent experiments. Error bars indicate the mean ± SD. **(G-J)** qPCR validation of four myogenic differentiation genes MyoD1, Myostatin, MYL, Myozenin that show upregulation upon G9a knockdown in RNA-Seq analysis. Data shown is representative of 2 independent experiments. Error bars indicate the mean ± SD. Statistical significance in **C-J** was calculated as unpaired two-tailed *t* test. \*\**P* ≤ 0.001, \*\*\**P* ≤ 0.001 and \*\*\*\**P* ≤ 0.0001.

**Supplemental Figure 4:**
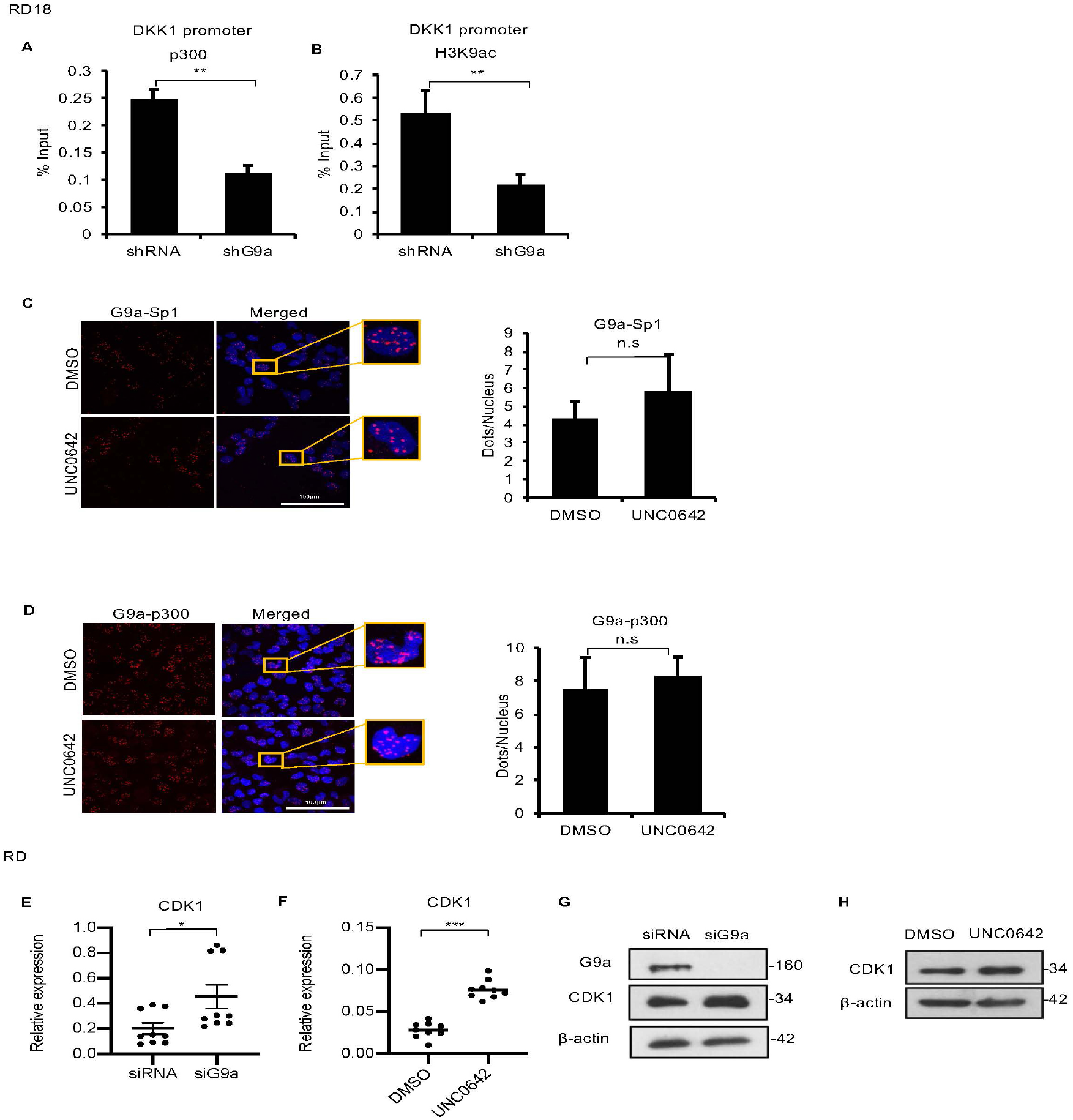
G9a regulates Sp1 and p300 occupancy at the *DKK1* promoter. **(A-B)** ChIP-PCR analysis showed decrease in p300 occupancy and H3K9ac enrichment at the *DKK1* promoter in shG9a RD cells as compared to control (n=2). Error bars indicate the mean ± SD. **(C)** PLA was used to determine G9a-Sp1 interaction which did not change upon UNC0642 treatment. Data shown is representative of 2 independent experiments. Error bars indicate the mean ± SD. **(D)** G9a-p300 interaction was not significantly altered by UNC0642 treatment. Data shown is representative of 2 independent experiments. Error bars indicate the mean ± SD. Statistical significance in **A, B, C and D** was calculated as unpaired two-tailed *t* test. \*\**P* ≤ 0.001, ns = not significant. **(E, F)** *CDK1* mRNA was analysed in control and siG9a RD cells and in control and UNC0642 treated RD cells (n=3). Values correspond to the average ± SEM. **(H, I)** CDK1 protein were analysed in in control and siG9a RD18 cells and in DMSO and UNC0642 treated RD18 cells. The image is representative of 2 independent experiments. Statistical significance in **E** and **F** was calculated by unpaired two-tailed *t* test. \*\*\**P* ≤ 0.001 and \*\*\*\**P* ≤ 0.0001.

